# Cubam receptor-mediated endocytosis in hindgut-derived pseudoplacenta of a viviparous teleost *Xenotoca eiseni*

**DOI:** 10.1101/2021.02.01.429082

**Authors:** Atsuo Iida, Kaori Sano, Mayu Inokuchi, Jumpei Nomura, Takayuki Suzuki, Mao Kuriki, Maina Sogabe, Daichi Susaki, Kaoru Tonosaki, Tetsu Kinoshita, Eiichi Hondo

**Affiliations:** Department of Animal Sciences, Graduate School of Bioagricultural Sciences, Nagoya University, Tokai National Higher Education and Research System, Nagoya, Aichi, Japan; Department of Chemistry, Faculty of Science, Josai University, Sakado, Saitama, Japan; Department of Aquatic Bioscience, Graduate School of Agricultural and Life Sciences, University of Tokyo, Bunkyo, Tokyo, Japan; Department of Regeneration Science and Engineering, Institute for Frontier Life and Medical Sciences, Kyoto University, Kyoto, Kyoto, Japan; Kihara Institute for Biological Research, Yokohama City University, Yokohama, Kanagawa, Japan

**Author notes:** **Correspondence**: Atsuo Iida, **E-mail**.

**Keywords:** endocytosis, Goodeidae, proteolysis, pseudoplacenta, teleost, viviparity

## Abstract

Nutrient transfer from mother to the embryo is essential for reproduction in viviparous animals. In the viviparous teleost *Xenotoca eiseni* belonging to the family Goodeidae, the intraovarian embryo intakes the maternal component secreted into the ovarian fluid via the trophotaenia. Our previous study reported that the epithelial layer cells of the trophotaenia incorporate a maternal protein via vesicle trafficking. However, the molecules responsible for the absorption were still elusive. Here, we focused on Cubam (<u>Cub</u>ilin-<u>Am</u>nionless) as a receptor involved in the absorption, and cathepsin L as a functional protease in the vesicles. Our results indicated that the Cubam receptor is distributed in the apical surface of the trophotaenia epithelium and then is taken into the intracellular vesicles. The trophotaenia possesses acidic organelles in epithelial layer cells and cathepsin L-dependent proteolysis activity. This evidence does not conflict with our hypothesis that receptor-mediated endocytosis and proteolysis play roles in maternal macromolecule absorption via the trohotaenia in viviparous teleosts. Such nutrient absorption involving endocytosis is not a specific trait in viviparous fish. Similar processes have been reported in the larval stage of oviparous fish or the suckling stage of viviparous mammals. Our findings suggest that the viviparous teleost acquired trophotaenia-based viviparity from a modification of the intestinal absorption system common in vertebrates. This is a fundamental study to understand the strategic variation of the reproductive system in vertebrates.

**Summary statement:** Here, we report that an endocytic pathway is a candidate for nutrient absorption in pseudoplacenta of a viviparous teleost. The trait may have developed from common intestinal mechanism among vertebrates.

## Introduction

August Krogh wrote, *“For such a large number of problems there will be some animal of choice or a few such animals on which it can be most conveniently studied”* [1]. This study aimed to investigate the molecular mechanism of maternal nutrient absorption in a species-specific pseudoplacenta of a viviparous teleost species belonging to the family Goodeidae.

Viviparity is a reproduction system, whereby the oocyte is fertilized in the female body, and subsequent embryo growth occurs with maternal component supply. Each viviparous animal has acquired processes specialized to the gestation in both the mother and embryo [2]. The placenta and umbilical cords in viviparous mammals are major components of the process for mother-to-embryo material transport [3,4]. Other viviparous components such as the extended yolk sac or pseudoplacenta are known in several viviparous vertebrates, except mammals [5].

The family Goodeidae is a freshwater small teleost distributed in Mexico, which includes approximately 40 viviparous species [6]. They possess trophotaenia, which is a hindgut-derived pseudoplacenta that absorbs the maternal component [7,8]. Trophotaenia is a ribbon-like structure consisting of a single epithelial layer, internal blood vessels, and connective tissues [9,10]. The epithelial cell is like an enterocyte in the intestine. Electron microscopy indicated that microvilli form in the apical side of the cell and intracellular vesicles in the cytoplasm [11]. Since the 1980s, these structures have been believed to be involved in maternal component absorption [12]. The nature of the maternal component was predicted to be proteins or other macromolecules secreted into the serum or ovarian fluids; however, no one has experimentally determined its distinct component [13,14]. Recently, we demonstrated that a yolk protein vitellogenin is secreted into the ovarian lumen of pregnant females, and the intraovarian embryo absorbs the nutrient protein via the trophotaenia using a goodeid species *Xenotoca eiseni* [15]. In that study, enterocyte-like microvilli and intracellular vesicles were also observed in the epithelial cells of the trophotaenia. We hypothesized that the epithelial layer cell in the trophotaenia absorbs the maternal protein and/or other components as macromolecules because the ovarian lumen lacks proteolysis activity like the digestive intestine. However, the molecules responsible for the trophotaenia-mediated macromolecule absorption have not been reported.

Vacuolated enterocytes involved in macromolecule absorption have also been reported in other vertebrate species, including suckling mammals and stomachless fish [16–18]. Park et al. [19] reported that the scavenger receptor complex Cubam (<u>Cub</u>ilin-<u>Am</u>nionless) and Dab2 are required for macromolecule uptake in zebrafish juveniles. Furthermore, the conditional knockout of *Dab2* in mice led to stunted growth and severe protein malnutrition at the suckling stage [19]. Based on this report, we discuss the commonality of molecular process for the macromolecule absorption between the intestinal enterocytes and the trophotaenia [15]. However, the molecules responsible for macromolecule absorption in the trophotaenia are still elusive.

Here, we report candidate molecules for nutrient uptake and subsequent proteolysis in the trophotaenia of *X. eiseni*. An RNA-Seq for trophoteniae indicated candidate receptor molecules and endocytosis-associated proteases expressed in the trophotaenia. Immunohistochemistry and biochemical assays suggested the presence and functions of the candidate factors in the trophotaenia.

## Methods

### Animal experiments

This study was approved by the ethics review board for animal experiments at Nagoya University (Approval number: AGR2020028). We sacrificed live animals in minimal numbers under anesthesia according to the institutional guidelines.

### Fish breeding

*X. eiseni* was purchased from Meito Suien Co., Ltd. (Nagoya, Japan). Adult fish were maintained in freshwater at 27 °C under a 14:10-h light: dark photoperiod cycle. Fish were bred in a mass-mating design, and approximately 30 adult fish were maintained for this study. The juveniles were fed live brine shrimp larvae, and the adults were fed Hikari Crest Micro Pellets and ultraviolet-sterilized frozen chironomid larvae (Kyorin Co., Ltd., Himeji, Japan). To accurately track the pregnancy period, the laboratory-born fish were crossed in a pair-mating design, and the mating behavior was recorded.

### Sample collection

Fish samples were anesthetized using tricaine on ice before the surgical extraction of tissues or embryos. The obtained samples were stored on ice until subsequent manipulations. In this study, we dissected approximately 10 pregnant females and extracted 15–30 embryos in each operation.

### RNA-Seq

Total RNA was extracted from trophotaenae of the 3^rd^ or 4^th^ week of the embryo extracted from the pregnant female using the RNeasy Plus Mini kit (QIAGEN). A total of six samples were obtained from three 3^rd^ week embryos and three 4^th^ week embryos. Next-generation sequencing (NGS) was outsourced to Macrogen Japan Corp. (Kyoto, Japan) using NovaSeq6000 (Illumina, Inc., San Diego, CA, USA). Approximately 60 million 150-bp paired-end reads were obtained in each sample. The NGS data was deposited to the DNA Data Bank of Japan (DDBJ, ID: DRA011209). *De novo assembly* and mapping to the reference sequence were performed by CLC Genomics Workbench (Filgen, Inc., Nagoya, Japan). The published transcript sequences of *Poecilia reticulata* (NCBI Genome, ID: 23338) were used as a reference. *P. reticulata* is one of the species related to *X. eiseni* for which the transcript sequences have been published, and it belongs to the order Cyprinodontiformes. The transcript sequences of *X. eiseni* were deposited into the DDBJ. The accession numbers are listed in Table 1.

**Table 1.**
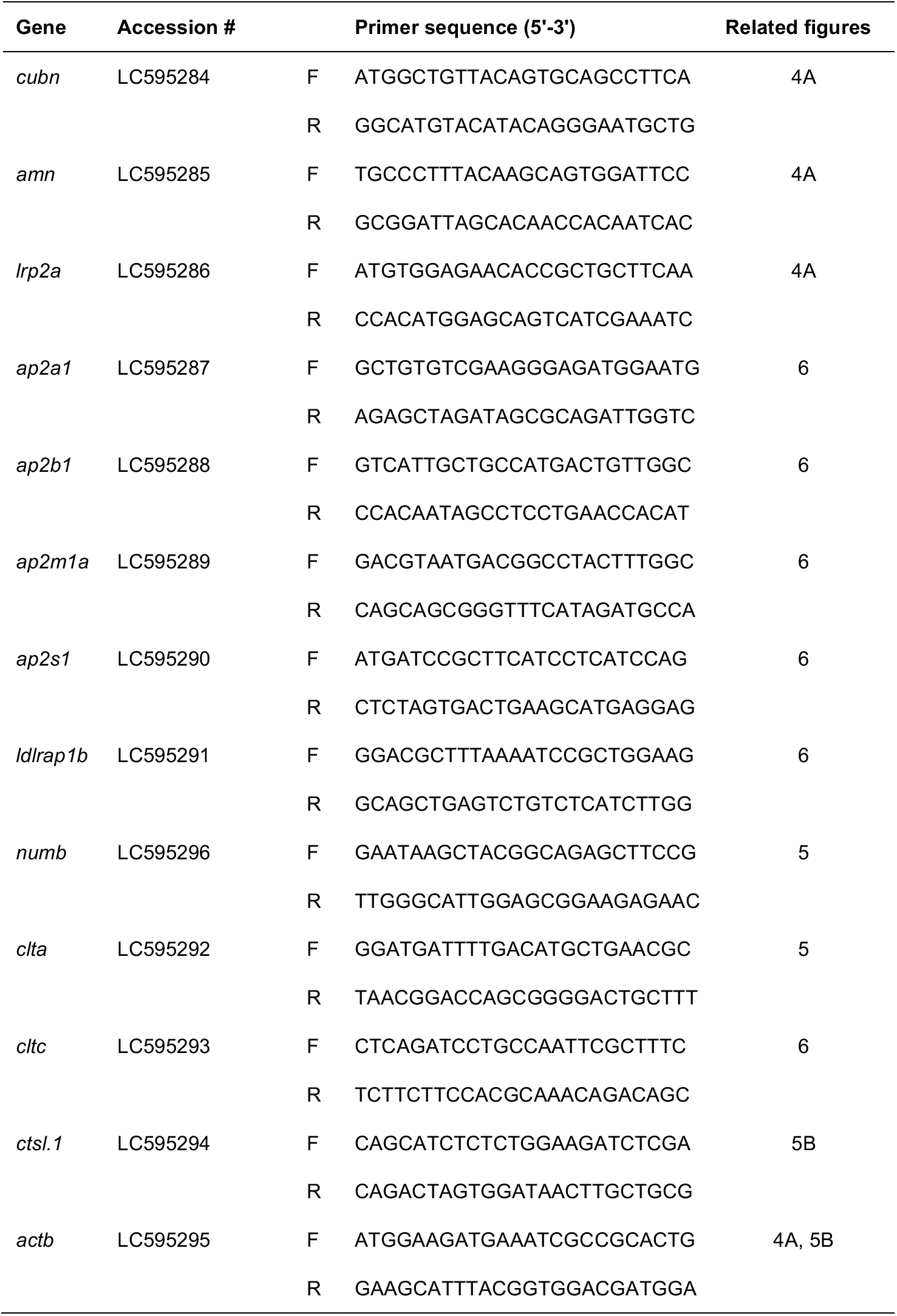
List of primers used in the study. F, forward, R, reverse.

### Phylogenetic analysis

Amino acid sequences for Cubn, Amn, or Cathepsin L of *Homo sapiens* (human), *Mus musculus* (house mouse), *Danio rerio* (zebrafish), *Oryzias latipes* (medakafish), *P. reticulata* (guppy), *Xiphophorus maculatus* (platyfish), and *X. eiseni* were collected from the NCBI protein database (https://www.ncbi.nlm.nih.gov/protein/). Phylogenetic trees were constructed by the neighbor-joining method with 1,000 bootstrap replicates in MEGAX (Ver.10.1.8) software (https://www.megasoftware.net/).

### Reverse transcription (RT) PCR

Total RNA was extracted from tissues or whole embryos using the RNeasy Plus Mini kit and reverse-transcribed using SuperScript IV reverse transcriptase (Thermo Fisher Scientific). PCR was carried out using KOD-FX-Neo (Toyobo, Osaka, Japan) under the following conditions: 100 s at 94 °C, followed by 32 cycles of 20 s at 94 °C, 20 s at 60 °C, 60 s at 72 °C, and 120 s at 72 °C. Primer sequences are listed in Table 1.

### Antibodies and antiserums

Antiserums against Cubn and Amn were generated in this study. The antigen sequences were 6× His-tagged peptide of 151 amino acids (aas) corresponding to the intermediate region of *X. eiseni* Cubn (Accession#, BCN28455; aa 691–841) and 6× His-tagged peptide of 247 aas corresponding to the C-terminal of *X. eiseni* Amn (Accession#, BCN28456). The experimental procedure has been described in our previous study [15]. All antibodies and antiserums used in this study are listed in Table S1.

### Immunohistochemistry

Tissue samples were fixed in 4.0% paraformaldehyde/phosphate-buffered saline (PFA/PBS) at 4 °C overnight. Samples were permeabilized using 0.5% TritonX-100/PBS at room temperature for 30 min. Endogenous peroxidase was inactivated by 3.0 % hydrogen peroxide/PBS for 10 min. Then, the sample was treated with Blocking-One solution (Nacalai Tesque, Kyoto, Japan) at room temperature for 1 h. Primary antibody or antiserums were used at 1:500 dilution with Blocking-One solution. Samples were incubated with primary antibody or antiserum at 4 °C overnight. Secondary antibodies were used at a 1:500 dilution in 0.1% Tween-20/PBS. Samples were treated with the secondary antibody solution at 4 °C for 2 h. We performed a 3,3’-Diaminobenzidine Tetrahydrochloride (DAB) color development using the DAB Peroxidase Substrate Kit, ImmPACT (Vector Laboratories, Inc., Burlingame, CA, USA). Microscopic observation was performed using an Olympus BX53 microscope and photographed using a DP25 digital camera (Olympus, Shinjuku, Japan).

### Fluorescent Immunohistochemistry

Tissue samples were fixed in 4.0% PFA/PBS at 4 °C overnight. Samples were permeabilized using 0.5% TritonX-100/PBS at room temperature for 30 min. Endogenous peroxidase was inactivated by 3.0 % hydrogen peroxide/PBS for 10 min. Then, the sample was treated with Blocking-One solution (Nacalai Tesque, Kyoto, Japan) at room temperature for 1 h. Primary antibody (anti-Cubn) was used at a 1:500 dilution with Blocking-One solution. Samples were incubated with primary antibody at 4 °C overnight. Secondary antibody was used at a 1:500 dilution in 0.1% Tween-20/PBS with 4’,6-diamidino-2-phenylindole (DAPI). Samples were treated with the secondary antibody solution at 4 °C overnight. Microscopic observation was performed using a Leica TCS SP8 microscope (Leica Microsystems, Wetzlar, Germany).

### Immunoelectron microscopy

Embryo samples were fixed in 4.0% PFA/PBS. Fixed samples were washed in PBS, then reacted with a primary antibody (anti-Cubn) at 4 °C overnight, and then reacted with biotinylated anti-rabbit IgG (Vector, Burlingame, CA, USA) at room temperature for 2 h. Samples were performed with the avidin–biotin–peroxidase complex kit (Vector), and visualized with 0.05% DAB (Dojindo Laboratories, Kumamoto, Japan) and 0.01% hydrogen peroxide in 50 mM Tris buffer (pH 7.2) at room temperature for 10 min. The procedure for electron microscopy is described in our previous study [15].

### Labeling acidic organelles

The live embryos immediately after extraction from the pregnant female at the 4^th^-week post-mating were incubated in PBS with a 1:1000 dilution of LysoTracker® Red (Thermo Fisher Scientific) for 1 h at room temperature. The samples were fixed with 4.0 % PFA/PBS and stained with DAPI. Microscopic observation was performed using a Leica DM5000 B microscope (Leica Microsystems, Wetzlar, Germany).

### Measurement of Cathepsin L activity

The trophotaenia lysate was prepared from six littermate embryos obtained from the pregnant females at the 4^th^-week post-mating. The trophotaeniae were manually extracted from the embryo under a microscope. Proteolysis detection was performed using the Cathepsin L Activity Fluorometric Assay Kit (BioVision, Inc., Milpitas, CA, USA). The fluorescence intensities were measured using a Qubit 4 Fluorometer (Thermo Fisher Scientific).

## Results

### Gene expression in trophotaenia

Our previous findings and the hypothesis based on that are described in Figure 1A. In a viviparous teleost, *X. eiseni*, the embryo is raised in the ovary while receiving maternal nutrients. The trophotaenia is a pseudoplacenta that plays a role in the absorption of maternal nutrients consisting of proteins and other supplements. Based on previous studies, we hypothesized that several maternal proteins are absorbed by endocytosis-mediated proteolysis in the epithelial cells of the trophotaenia. To verify this hypothesis, we explored candidate genes for receptor, adaptor, vesicle, and protease proteins that are highly expressed in the trophotaenia. RNA-Seq analyses were performed using total RNA extracted from the trophotaeniae of the 3rd- or 4th-week embryos (Figure 1B). We selected candidate genes and compared their predicted expression level as transcription per million (TPM) values that were calculated using the known transcript sequence of *P. reticulata* as reference. RNA-Seq suggested that *cubilin (cubn*) and *amnionless (amn*) were highly expressed in the trophotaeniae; however, other co-receptor genes (*lrp1aa, lrp2a*) were considerably lower (Figure 1C, Table 2). Adaptor protein-2 (AP2) subunit genes (*ap2a1, ap2b1, ap2m1a*, and *ap2s1*) were more highly expressed than the other family adaptor genes (*ldlrap1b* and *numb*) (Figure 1C, Table 2). Two families of the vesicle coating protein genes (*clta*, *cltbb, cltc; flot1b, flot2a*) were expressed higher than the vesicle proteins classified in other families (*cav2* and *cav3*) (Figure 1C, Table 2). The lysosomal endopeptidase enzyme gene (*ctsl.1*) exhibited a high TPM value (> 10,000) rather than that of not only protease family genes but also most of the genes expressed in the trophotaeniae (Figure 1C, Table 2). In this investigation, we assumed that the 3^rd^ week is a good growth period (mid pregnancy) and that the 4^th^ week is a poorer growth (late pregnancy) period; thus, we expected that the transcriptomes related to the nutrient transport would vary between these stages. However, the trend of the predicted TPM values in most of the selected genes was not noticeably different between the samples. In the 4^th^ week, only one *clathrin* and four *cathepsin genes* were significantly higher than levels in the 3^rd^ week (Figure 1C).

**Figure 1.**
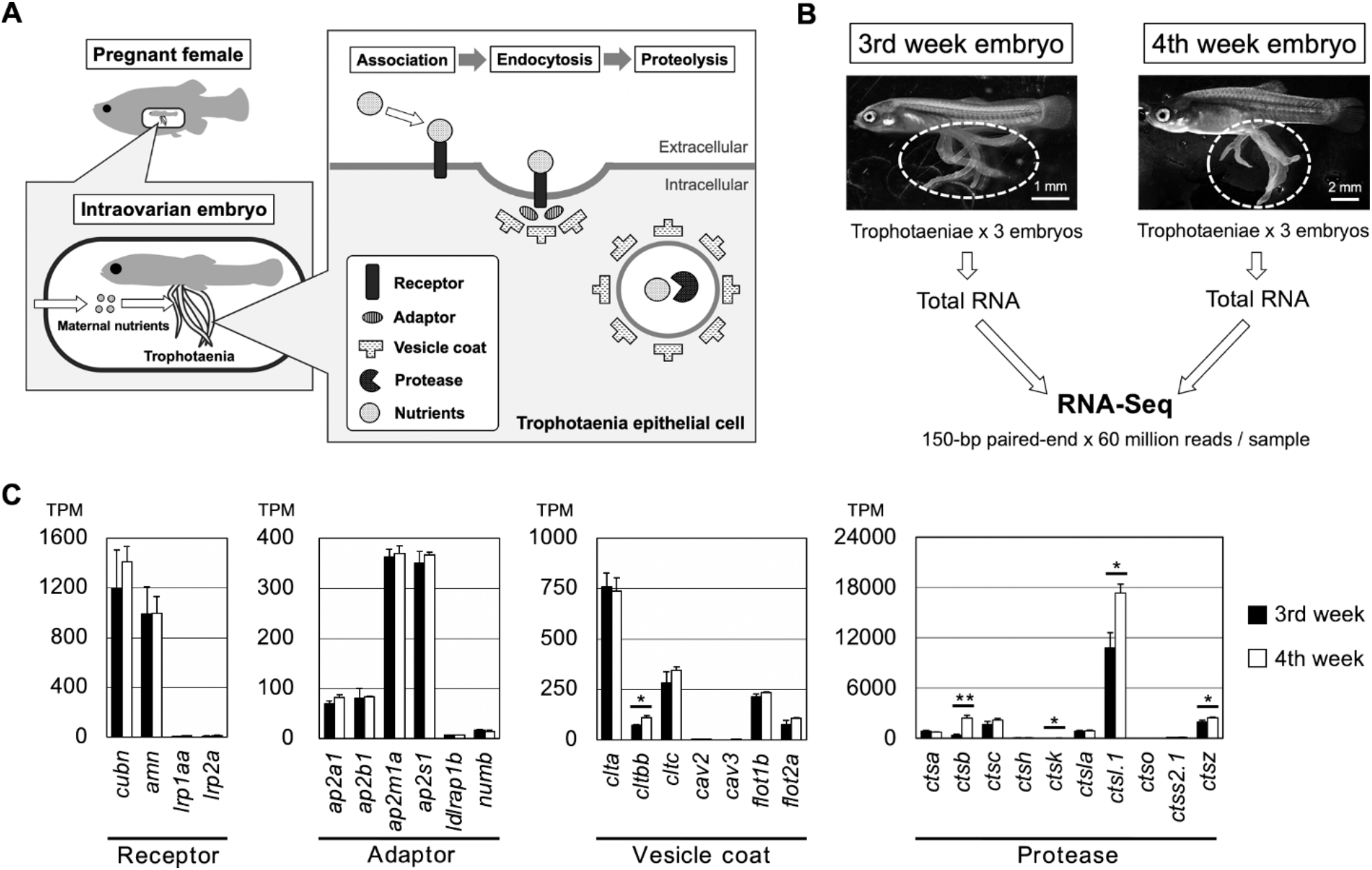
Exploring candidate genes for nutrient absorption. **A**. A working model and hypothesis of this study. In the goodeid viviparous fish (*X. eiseni*), intraovarian embryo absorbs maternal nutrients via the trophotaenia. We hypothesized that endocytosis-mediated proteolysis is related to nutrient absorption. Based on the scenario, potential target genes are for endocytic receptors, adaptors, vesicle coating proteins, and proteases. **B**. An experimental scheme for the RNA-Seq analysis. RNA samples were obtained from the trophotaeniae (white dotted line) of the single intraovarian embryos extracted from the pregnant females of the 3^rd^- or 4^th^-week post-mating. The RNA-Seq was performed using three samples at every stage. **C**. The graphs indicate the transcript per million (TPM) values of the genes selected from the RNA-Seq data that are involved in the endocytosis-mediated proteolysis pathway. Student’s *t* test was used for statistical analyses. * *p* < 0.05. ** *p* < 0.01. Error bars represent the standard error.

**Table 2.**
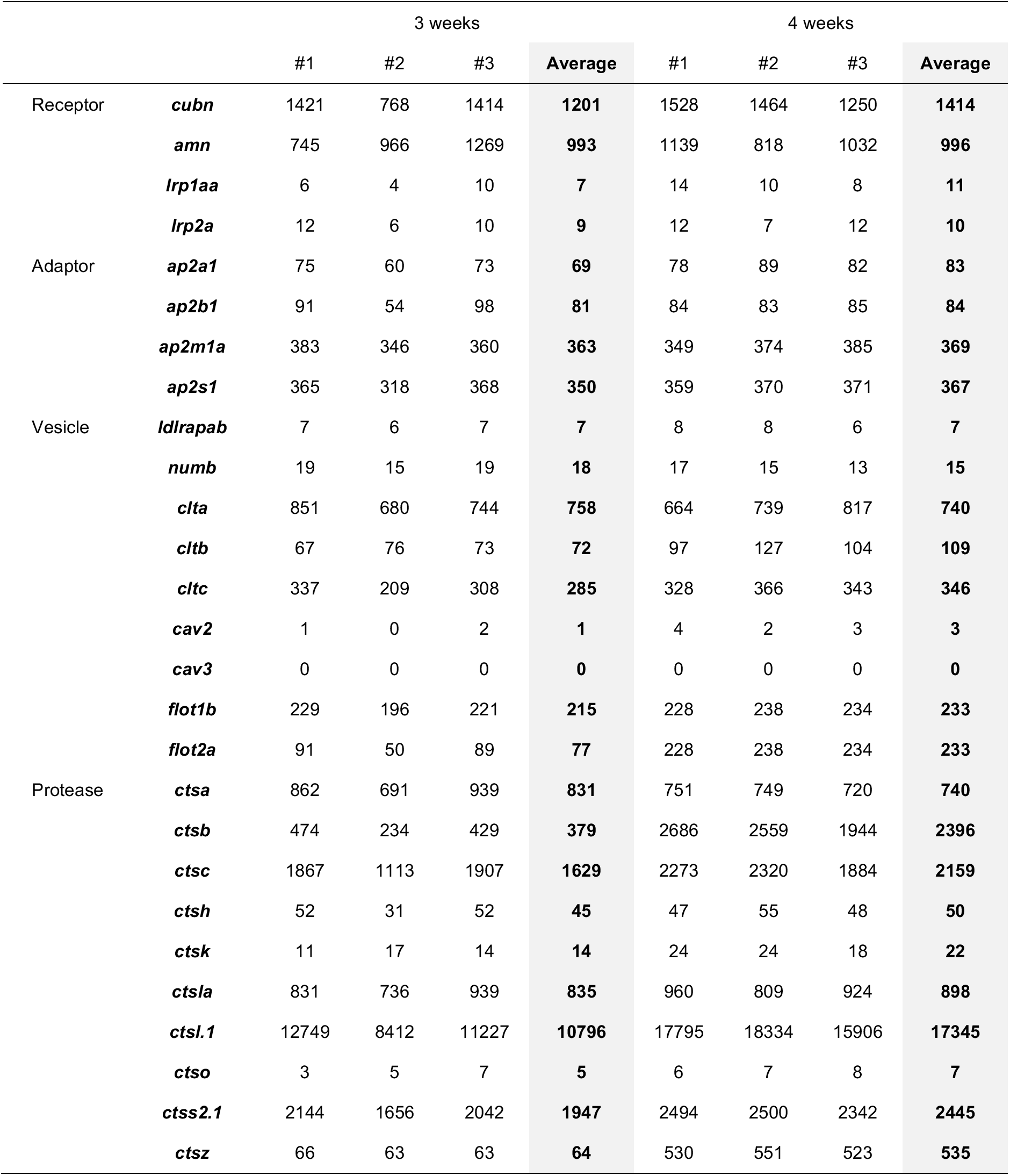
The TPM values for endocytic genes in the RNA-seq analysis.

### Sequences for Cubam receptor genes

A Cubam receptor is known to be a membrane-bound multi-ligand receptor consisting of Cubn and Amn (Figure 2A). The secreted protein Cubn specifically binds to the transmembrane protein Amn; thus, the CUB domain that associates with the ligands are localized on the apical surface of the plasma membrane. We obtained the amino acid sequences of *X. eiseni* Cubn and Amn by *de novo* assembly of NGS reads from the trophotaeniae. The assembled sequences of *X. eiseni* were calculated to be close to the orthologous sequences of *P. reticulata* and *X. maculatus* (Figure 3A and B). The sequences of Cubn and Amn were compared among four vertebrate species, *H. sapiens*, *D. rerio*, *P. reticulata*, and *X. eiseni*. The binding motifs were conserved among the species (Figure 2B and C). A phenylalanine-any-asparagine-proline-any-phenylalanine (<u>F</u>X<u>NP</u>X<u>F</u>) amino acid sequence in the intracellular domain of Amn is known to be a conserved motif used to bind adaptor proteins. The motifs were also conserved among the species (Figure 2D).

**Figure 2.**
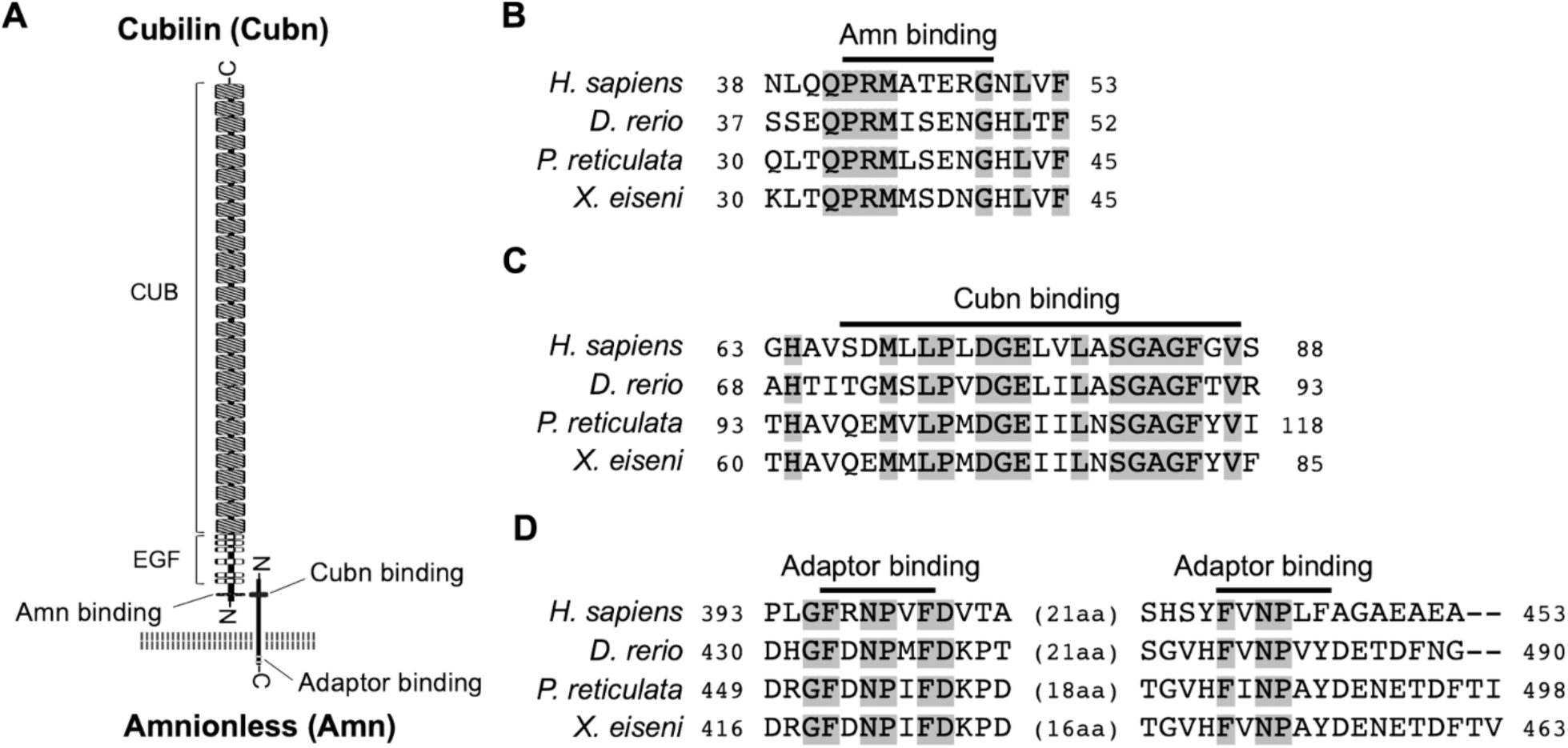
Structures and amino acid sequences for *X. eiseni* Cubn and Amn. **A**. The illustration indicates that a typical structure of Cubam receptor complex consists of Cubn and Amn. Both proteins possess a conserved motif to bind each other in the N-terminal regions, and the Amn possesses two adaptor binding motifs in the C-terminal intracellular domain. **B-D**. A comparison of amino acid sequence around the Amn-binding motif in Cubn (**B**), Cubn-binding motif in Amn (**C**), and adaptor binding motifs in Amn (**D**) between *X. eiseni* and three other vertebrate species. The gray filled texts are conserved sequences among the four species.

**Figure 3.**
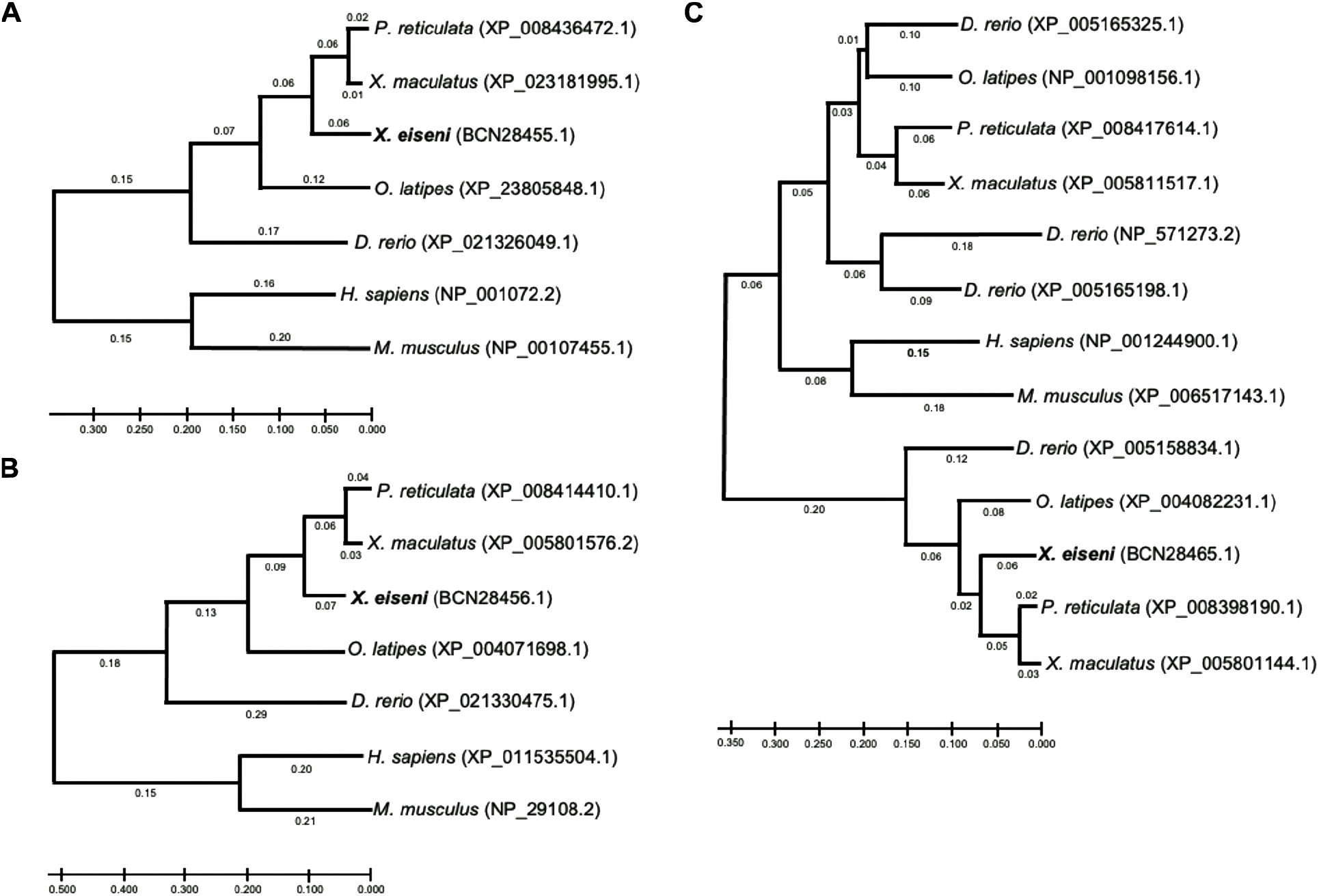
Phylogenetic comparison of Cubn, Amn, and Cathepsin L amino acid sequences. Phylogenetic tree indicating the relationship among the amino acid sequences of Cubn (**A**), Amn (**B**), or Cathepsin L (**C**) in the seven vertebrate species. The sequences of *X. eiseni* (Cyprinodontiformes: Goodeidae) identified in this study were calculated to be closed to the sequences of orthologues in *P. reticulata* and *X. maculatus* (Cyprinodontiformes: Poeciliidae).

### Distribution of Cubilin and Amnionnless in trophotaenia

To validate the expression of *cubn* and *amn* in the trophotaenia, semi-quantitative RT-PCR analyses were performed using total RNAs extracted from the whole-embryo including the trophotaeniae, isolated trophotaeniae, and adult skeletal muscle. The muscle sample was used as a control for tissue with low endocytosis activity. The gene expression patterns did not conflict between the results of RT-PCR and RNA-Seq. In the embryo and trophotaenia, both *cubn* and *amn* were more highly expressed than *lrp2a* (Figure 4A). Conversely, no or few expressions of the target genes except *actb* as a positive control were detected in the adult muscle. Next, to detect protein localization, immunohistochemistry was performed using antibodies against Cubn or Amn. In both proteins, strong signals were observed in the epithelial monolayer of the trophotaenia, while few background noises were detected in the control assay (Figure 4B-F, see also Figure S1 and S2). Confocal microscopy revealed the cellular distribution of the anti-Cubn signals. The signals in the apical surface of the epithelial cell were detected as a homogeneous distribution, while almost all signals in the cytoplasm were captured as a dot pattern (Figure 4G, Figure S3). Immunoelectron microscopy revealed that the microvilli were distributed on the apical surface of the trophotaenia epithelium, and intracellular vesicles were observed in the cytoplasm; furthermore, anti-Cubn signals were distributed in the intracellular vesicles and overlapped with the microvilli on the apical surface (Figure 4H).

**Figure 4.**
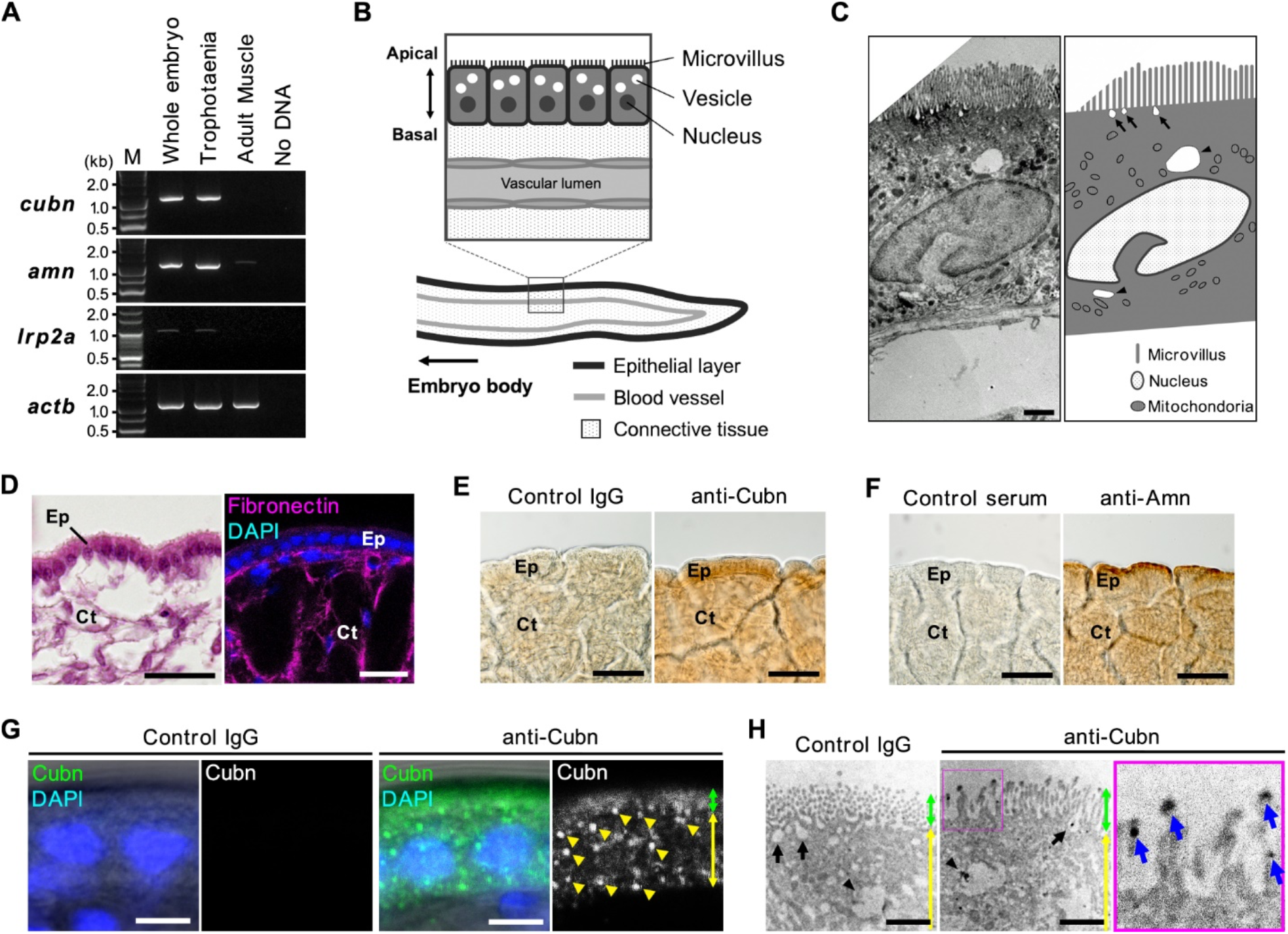
Distribution of receptors involved in endocytosis. **A**. RT-PCR for the candidate genes for the receptors involved in the endocytosis. All amplicons were detected as single band on the expected sizes based on the transcript sequences obtained from the *de novo* assembly. **B**. The illustration indicates an internal structure of the trophotaenia. An epithelium cell layer configures an outermost structure of the trophotaenia, which contacts the ovarian luminal fluids. The layer consists of an enterocyte-like cell with microvilli on the apical surface and vesicles in the cytoplasm. **C**. Electron microscopy (left) and corresponding illustration (right) of an epithelial layer cell of the trophotaenia. Microvilli are distributed on the apical surface of the cells. Arrows indicate endocytic vesicles during the invagination phase. Arrowheads indicate intracellular vesicles after endocytosis. Scale bar: 1 μm. **D**. The structure of the trophotaeniae stained with HE (left) or fluorescent immunohistochemistry. A fibronectin antibody was used to visualize the connective tissues inside the epithelial layer. Ep, epithelial layer. Ct, connective tissue. Scale bar, 25 μm. **E-F**. Immunohistochemistry using the Cubn antibody or Amn antiserum in the trophotaenia. In both samples, DAB color development was observed in the epithelial layer. Ep, epithelial layer. Ct, connective tissue. Scale bar, 50 μm. **G**. Confocal microscopy images of fluorescent immunohistochemistry using the Cubn antibody in the epithelial layer cells. Yellow triangles indicate the dotted signals in the epithelial cells of the trophotaenia. Green double-headed arrow indicates the apical surface defined by the flat signal. Yellow double-headed arrow indicates the cytoplasmic region including the dotted signals. Scale bar, 5 μm. **H**. Immunoelectron microscopy using the Cubn antibody in the epithelial layer cells. Green double-headed arrow indicates the microvilli on the apical surface. Yellow double-headed arrow indicates the cytoplasmic region including the dotted signals. Arrows indicate endocytic vesicles in the invagination phase. Arrowheads indicate intracellular vesicles after endocytosis. Anti-Cubilin signals were observed in both vesicles. The enlarged image is corresponding to the magenta square focused on the microvilli. Blue arrows indicate anti-Cubn signals in the microvilli. Scale bar, 1 μm.

### Proteolysis activity in trophotaenia

Cathepsin L is a lysosomal cysteine proteinase characterized by three conserved protease regions and active sites consisting of cysteine, histidine, or asparagine (Figure 5A). The functional regions were conserved in the *X. eiseni* Cathepsin L protein translated from the coding sequence of *ctsl.1*. The whole amino acid sequence of *X. eiseni* Cathepsin L was calculated to be close to the orthologous proteins of *P. reticulata* and *X. maculatus* (Figure 3C). RT-PCR analysis revealed strong expression of *ctsl.1* in trophotaenia (Figure 5B). To identify the type of cells that have proteolysis activity in trophotaenia, acidic organelles including lysosomes and endosomes were detected using a fluorescent probe. LysoTracker® analysis indicated the presence of acidic organelles in the epithelial layer cells (Figure 5C). The signals were distributed in the cytoplasm and were not components of the nuclei (Figure 5D). According to the RNA-Seq analysis, *ctsl.1* was presumed to be the highest expressed *cathepsin* gene in trophotaenia; thus, we calibrated the proteolysis activity of cathepsin L in trophotaenia using a fluorescent substrate-based measurement system. The fluorescence, indicating substrate digestion, was significantly higher in the trophotaenia lysate than in the control at 1 h after reagent mixing. Furthermore, the intensity for the lysate continued to increase for 7 h (Figure 5E, Table 3). Conversely, the increase in intensity was strongly suppressed by a cathepsin L inhibitor. The fluorescent values at each timepoint were statistically different from each other.

**Figure 5.**
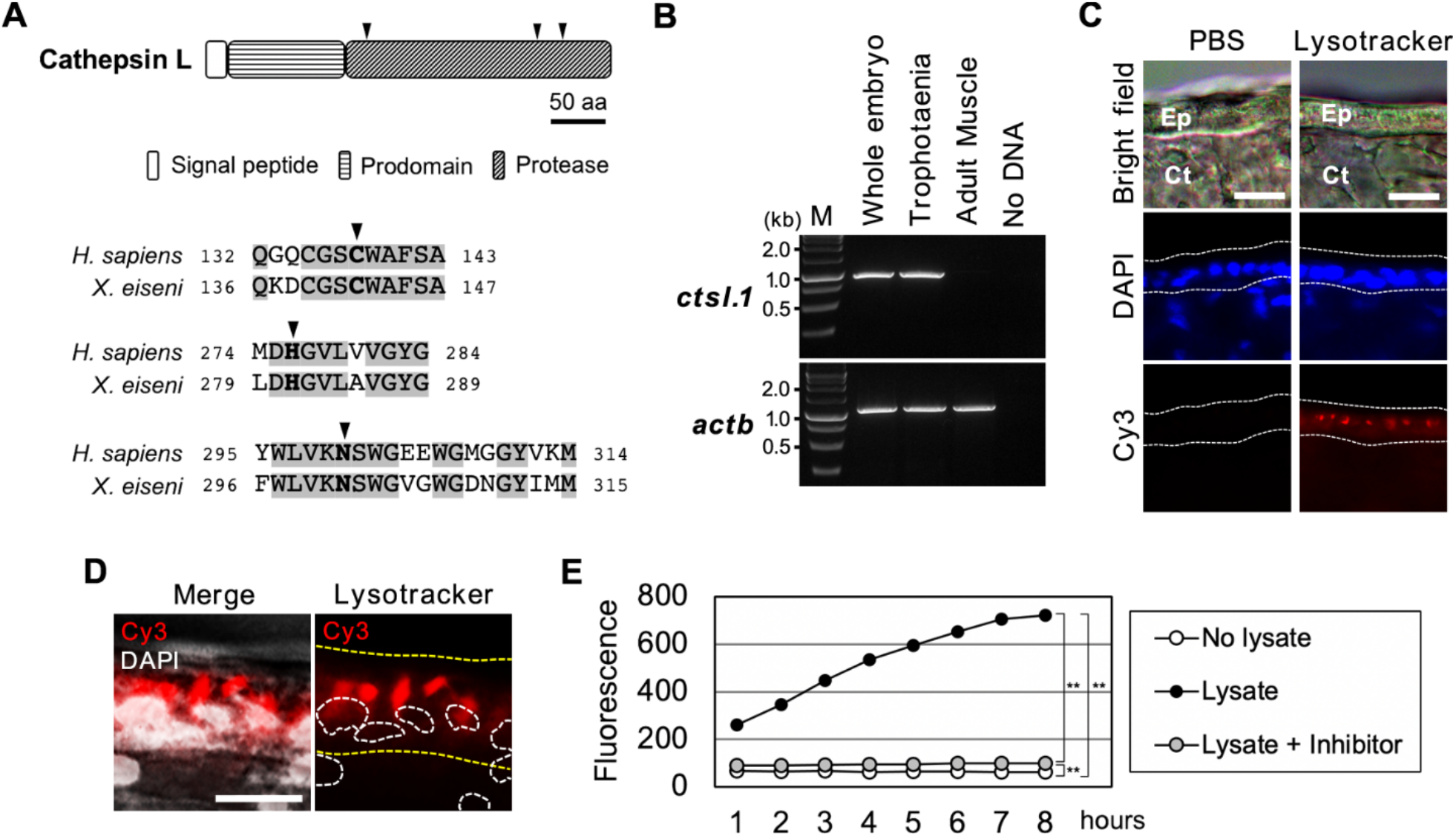
Proteolysis activities in trophotaenia. **A**. The illustrations indicate a typical structure of cathepsin L and a comparison of the protease domains of cathepsin L between *H. sapiens* and *X. eiseni*. The gray filled texts are conserved sequences among the species. The black triangles indicate protease active sites. **B**. RT-PCR for *cstl.1* exhibited the highest TPM value in the RNA-Seq analysis. The amplicons were detected as a single band on the expected size based on the transcript sequences obtained from the *de novo* assembly. **C**. Labeling of acidic organelles including lysosomal vesicle in the trophotaenia. The lysotracker treatment exhibited red fluorescence in the epithelial layer (white dotted line). Ep, epithelial layer. Ct, connective tissue. Scale bar, 20 μm. **D**. High magnification image of the lysotracker-treated epithelial cell layer of the trophotaenia. Yellow dotted line indicates the epithelial layer and white dotted circles are the nuclei of the epithelial cells. The lysotracker fluorescence was observed in the cytoplasm. Scale bar, 10 μm. **E**. Measurement of cathepsin L activity based on fluorescent substrate degradation. The vertical line indicates the accumulation of cleaved fluorescent substrates, which means cathepsin L activity in the sample solution. The fluorescent value of the trophotaenia lysate sample was increased by 8 h after the reaction started. Student’s *t* test was used for statistical analyses. ** *p* < 0.01.

**Table 3.** The measured fluorescent signal values indicating cathepsin L-dependent proteolysis.

|  |  | 1 h | 2 h | 3 h | 4 h | 5 h | 6 h | 7 h | 8 h |
| --- | --- | --- | --- | --- | --- | --- | --- | --- | --- |
| **No lysate** | #1 | 68.8 | 67.8 | 67.2 | 65.8 | 65.5 | 65.8 | 65.6 | 65.2 |
|  | #2 | 66.5 | 65.9 | 64.8 | 62.7 | 64.0 | 64.4 | 63.8 | 62.7 |
|  | #3 | 66.8 | 65.4 | 69.8 | 63.1 | 63.3 | 62.8 | 61.8 | 62.6 |
|  | **Average** | **67.4** | **66.3** | **67.3** | **63.8** | **64.3** | **64.3** | **63.8** | **63.5** |
| **Lysate** | #1 | 260.3 | 345.0 | 437.7 | 530.4 | 579.7 | 637.3 | 699.6 | 694.1 |
|  | #2 | 264.9 | 351.6 | 460.3 | 551.3 | 617.0 | 678.4 | 727.0 | 748.2 |
|  | #3 | 256.6 | 339.2 | 441.8 | 522.7 | 587.1 | 642.9 | 692.5 | 720.8 |
|  | **Average** | **260.6** | **345.3** | **446.6** | **534.8** | **594.6** | **652.9** | **706.3** | **721.0** |
| **Lysate + inhibitor** | #1 | 94.0 | 96.2 | 98.0 | 99.6 | 101.0 | 106.8 | 104.4 | 101.9 |
|  | #2 | 88.1 | 88.8 | 91.1 | 91.5 | 93.4 | 94.0 | 98.0 | 98.2 |
|  | #3 | 88.3 | 88.4 | 89.4 | 93.1 | 94.2 | 96.4 | 99.4 | 101.0 |
|  | **Average** | **90.1** | **91.2** | **92.8** | **94.7** | **96.2** | **99.0** | **100.6** | **100.4** |

### Adaptors and vesicle coating proteins

The expression of candidate genes for adaptors (*ap2a1, ap2b1, ap2m1a, ap2s1, ldlrap1b*, and *numb*) and vesicle coating proteins (*clta* and *cltc*) were confirmed by RT-PCR (Figure 6). We determined an incomplete transcript for *X. eiseni* Dab2 from the *de novo* assembly; however, it lacked internal sequences in comparison with the proteins in other vertebrates (Figure S4). Furthermore, no amplicons were obtained by RT-PCR using primers designed based on the Dab2-like sequence.

**Figure 6.**
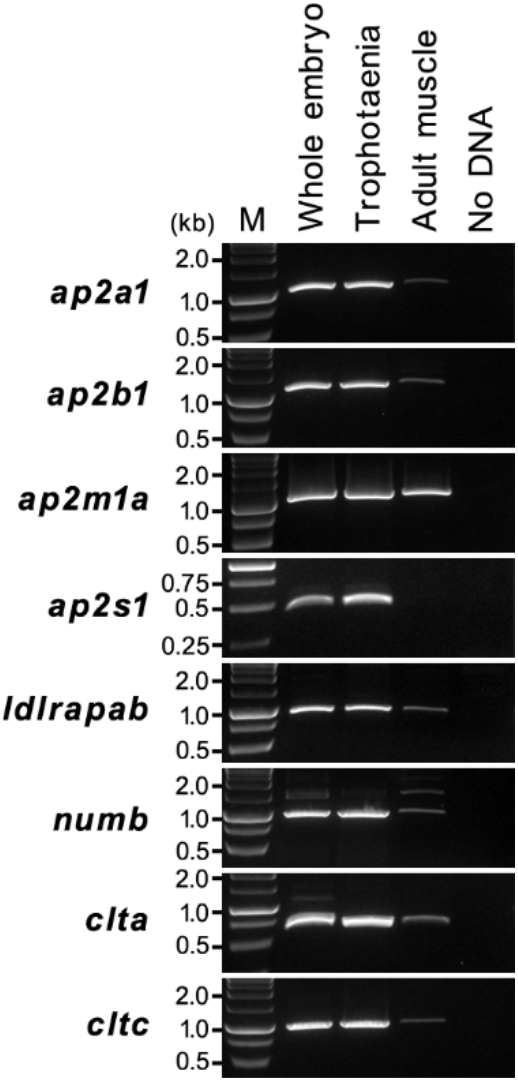
Gene expression analysis of the endocytic adaptor and vesicle coating genes. RT-PCR was performed for the candidate genes encoding the adaptors and vesicle coating genes involved in endocytosis. The amplicons were detected as single bands of the expected sizes based on the transcript sequences obtained from the *de novo* assembly. M, marker.

## Discussion

RNA-Seq analyses revealed high expression of the genes for receptor-mediated endocytosis in the trophotaenia of *X. eiseni* embryos. Cubn and Amn form a Cubam receptor complex that associates with Vitamin-B12, albumin, transferrin, and other ligands [20,21]. The predicted amino acid sequences for *X. eiseni* Cubn and Amn obtained from the *de novo* assembly both possessed a conserved motif which allowed binding to each other, and Amn retained adaptor binding sites in the intracellular region [21, 22]. Immunohistochemical analysis indicated the presence of Cubn not only in the apical surface of the epithelial layer but also in the intracellular vesicles in the cells, suggesting the incorporation and recycling of endocytic receptors [23,24]. This evidence supports the idea that Cubam plays a role as a receptor for the intraovarian nutrients and is involved in endocytosis. We also indicated the low presence of Lrp2 (also known Megalin), which is a major co-receptor for Cubam [25]. This indicated that Cubam works independently or in cooperation with other co-receptors, except Lrp2, in the trophotaenia. In zebrafish, a previous study reported that Cubam-dependent endocytosis in the lysosome rich enterocytes (LREs) does not require the presence of Lrp2 [19]. This supports our hypothesis; however, we do not exclude the possibility that our sequencing and alignment processes could not catch other *lrp2* orthologous genes in *X. eiseni*. Park et al. [19] also reported that Dab2 is an essential adaptor molecule not only in the zebrafish larval intestine but also in the endocytic nutrient absorption in the ileum of suckling mice. However, we obtained only an incomplete sequence for the *X. eiseni* Dab2-like protein without an internal region compared with mammalian orthologs. This may be an alternative splice form with less endocytic activity [26,27]. Furthermore, we did not detect Dab2-like expression by RT-PCR, and we could not exclude the possibility that the predicted Dab2-like sequence is caused by an assembly error. Thus, the contribution of Dab2 to endocytosis in the trophotaenia has not been confirmed in this study. The other adaptor or vesicle coating proteins we detected were typical co-factors for receptor-mediated endocytosis. Conversely, Caveolin is a vesicle coating protein involved in receptor-independent endocytosis [28]. Thus, a low expression of *caveolin* genes (*cav2* and *cav3*) does not conflict with the activation of Cubam-mediated endocytosis. Additionally, biochemical assays supported that cathepsin L is an active protease that functions in the intracellular vesicles that is configured following membrane budding. This evidence supports the idea that Cubam-mediated endocytosis and cathepsin L-dependent proteolysis is one of the key mechanisms for the absorption of the maternal component in *X. eiseni* embryos.

Cubam is also known to be a scavenger receptor involved in nonspecific protein uptake [20]. In this study, we did not determine the Cubam ligands in the trophotaenia. However, the candidates are limited to intraovarian secreted proteins. In several goodeid species, previous studies indicated that the ovarian fluids include various proteins in a pattern similar to that in the blood serum [13,29]. Therefore, several serum proteins, albumen, transferrin, and others known as Cubam ligands, are potential targets in the case of intraovarian nutrient absorption via the trophotaenia. Another possibility is that vitellogenin is also a potential target because it is secreted into the ovarian fluids and is absorbed into the trophotaenia via intracellular vesicles [15]. Another study suggested that the Cubn-Amn-Lrp2 receptor complex is related to transport of yolk proteins, including vitellogenin, in endodermal epithelial cells during chicken yolk sac growth [30]. These reports support the idea that Cubn and Amn are potent candidate molecules for a receptor complex involved in maternal component uptake in the trophotaenia. However, functional analyses such as gene knockout or transgenic technologies used in conventional model animals, such as laboratory mice or small oviparous fish, have not been applied to goodeid fish. For a direct validation of our hypothesis, these reverse genetic methods should also be developed in the viviparous teleost.

As described above, endocytosis-mediated nutrient absorption is not limited to the pseudoplacenta of the viviparous fish; it has also been reported in intestinal tissues (Figure 7). In mammals, macromolecule absorption in the intestine is limited to the suckling period [31]. In the stomachless fish, endocytosis-mediated absorption is considered to continue in part of the intestine for life, because of low digestive activity in the enteric cavity [18]. We found that Cubn and Amn are also distributed in the intestinal epithelial cells of adult *X. eiseni*. (Figure S5). In most invertebrate species, because the extracellular digestive system is primitive, food particles are absorbed into the intestinal cells by vesicle trafficking and degraded by intracellular digestion [32, 33]. Thus, we hypothesize that endocytic nutrient absorption and intracellular digestion are ancestral traits common not only in vertebrates but also in invertebrates, and their importance has decreased in certain vertebrates with development of the extracellular digestive system in the enteric cavity. The ancestor of the goodeid species may have applied the endocytic process to the reproductive system, and then configured the unique hind-gut derived pseudoplacenta for matrotrophic nutrition during gestation [7,34]. To validate our hypothesis, further exhaustive omics analyses using the goodeid species and comparative research using the subfamily Empetrichthyinae, which is the oviparous species most closely related to the viviparous goodeid fish, are required [6,35].

**Figure 7.**
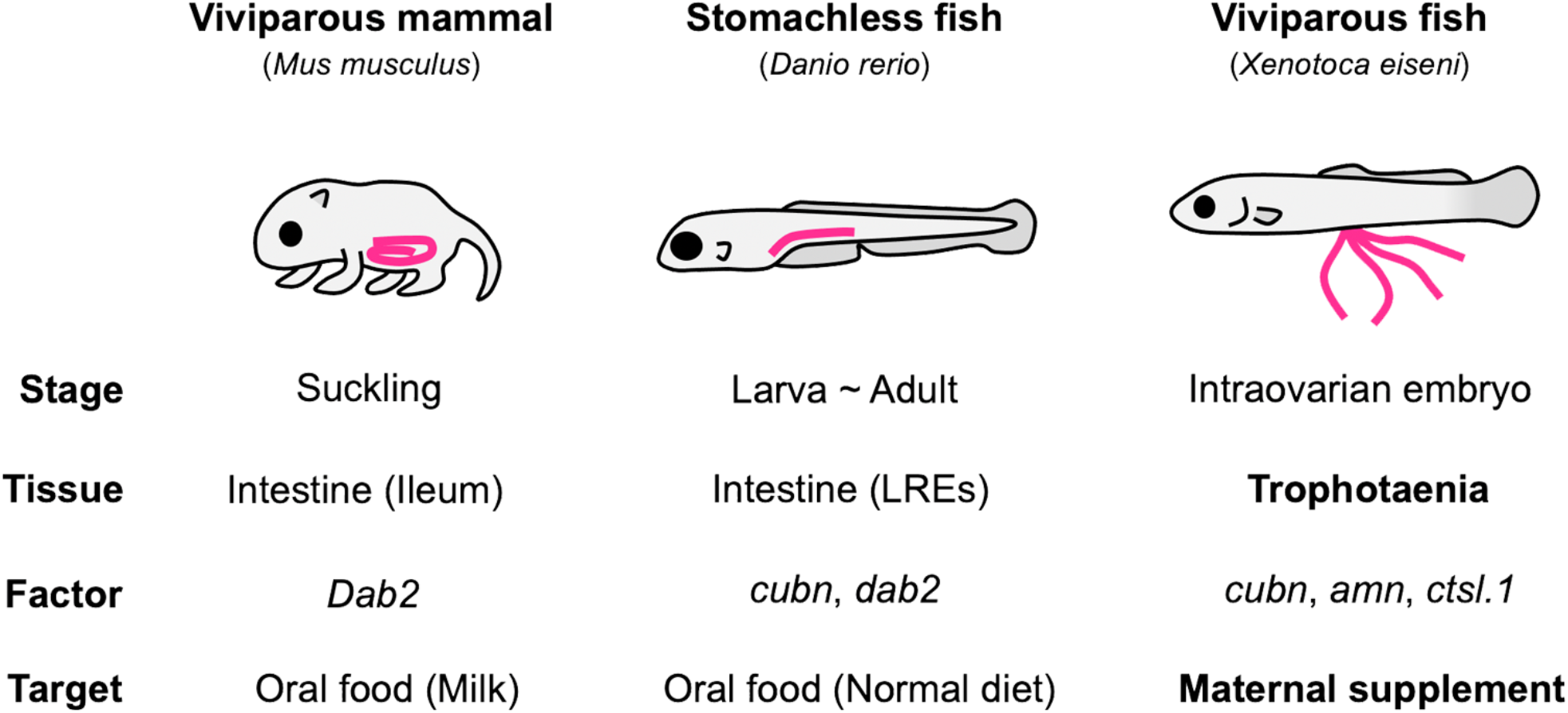
Comparison of endocytosis-mediated nutrient intake in vertebrates. Endocytosis-mediated nutrient intake has also been reported in a viviparous mammal and stomachless fish. In these species, oral ingestion macromolecules are absorbed from a region of the intestine via endocytosis. In contrast, intraovarian embryos of the viviparous fish (family Goodeidae) absorbs the maternal component in the ovarian fluids from the trophotaeniae via endocytosis. Endocytosis in each species is predicted to be driven by similar molecular process in the enterocytes of the intestine or the enterocyte-like cells of trophotaenia; however, the biological ontology would be divergent between the viviparous fish (intraovarian growth) and the others (feeding after birth).

This study is an investigation of species-specific traits based on the transcriptome of a non-conventional model species. The results revealed potential candidate molecules for nutrient absorption in the pseudoplacenta of the viviparous teleost. As August Krogh wrote, this kind of study would be suitable for investigation using the most appropriate species and is unsuitable for verification using alternative models such as viviparous rodents or oviparous teleosts. We believe that this study is an important and fundamental step in understanding the strategic variation of the reproductive system in vertebrates.

## Acknowledgments

This work was supported by research grants from the Nakatsuji Foresight Foundation and the Daiko Foundation.

## Competing interest statement

The authors declare that they have no competing interests.

## Data accessibility

The data that support the findings have been provided with the manuscript.

## Notes

### Competing Interest Statement

The authors have declared no competing interest.

