## Supplementary material for "Cubam receptor-mediated endocytosis in hindgut-derived pseudoplacenta of a viviparous teleost *Xenotoca eiseni*": SI for JEB.pdf (from another version of this preprint: version 4, 2021-03-22)

**This PDF file includes:**

Figures S1 to S5

Tables S1

**Additional Figures**

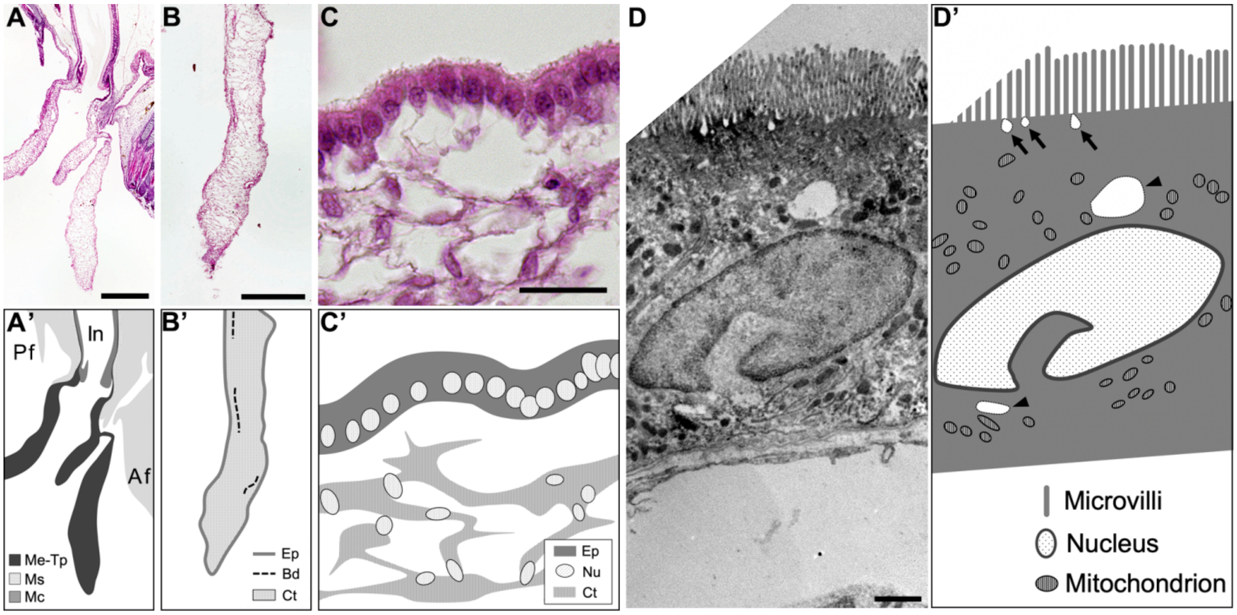


**Figure S1. Tissue structures of the trophotaenia in *Xenotoca eiseni* embryos.**

**A-A'**. Hematoxylin and eosin (HE)-stained section (A) and corresponding illustration (A') of the tissue structure around the anus at the border of the hindgut and trophotaenia of the embryo at the fourth gestational week. Trophotaeniae are sequential to the mesenchymal layer of the intestine. In, intestine; Pf, pelvic fin; Af, anal fin; Me-Tp, mesenchyme-to-trophotaenia; Ms, muscle layer; Mc, mucosal layer. Scale bar: 500 µm. **B-B'**. HE-stained section (B) and corresponding illustration (B') of the internal structure of the trophotaenia. Ep, epithelial layer cells; Bd, blood vessel; Ct, connective tissue. Scale bar: 500 µm. **C-C'**. Enlarged image of the HE-stained section (C) and corresponding illustration (C') of the epithelial layer of the trophotaenia. Ep, epithelial layer; Nu, nucleus; Ct, connective tissue. Scale bar: 20 µm. **D-D'**. Electron microscopy (D) and corresponding illustration (D') of an epithelial layer cell of the trophotaenia. Microvilli are distributed on the apical surface of the cells. Arrows indicate endocytic vesicles during the invagination phase. Arrowheads indicate intracellular vesicles after endocytosis. Scale bar: 1 μm.

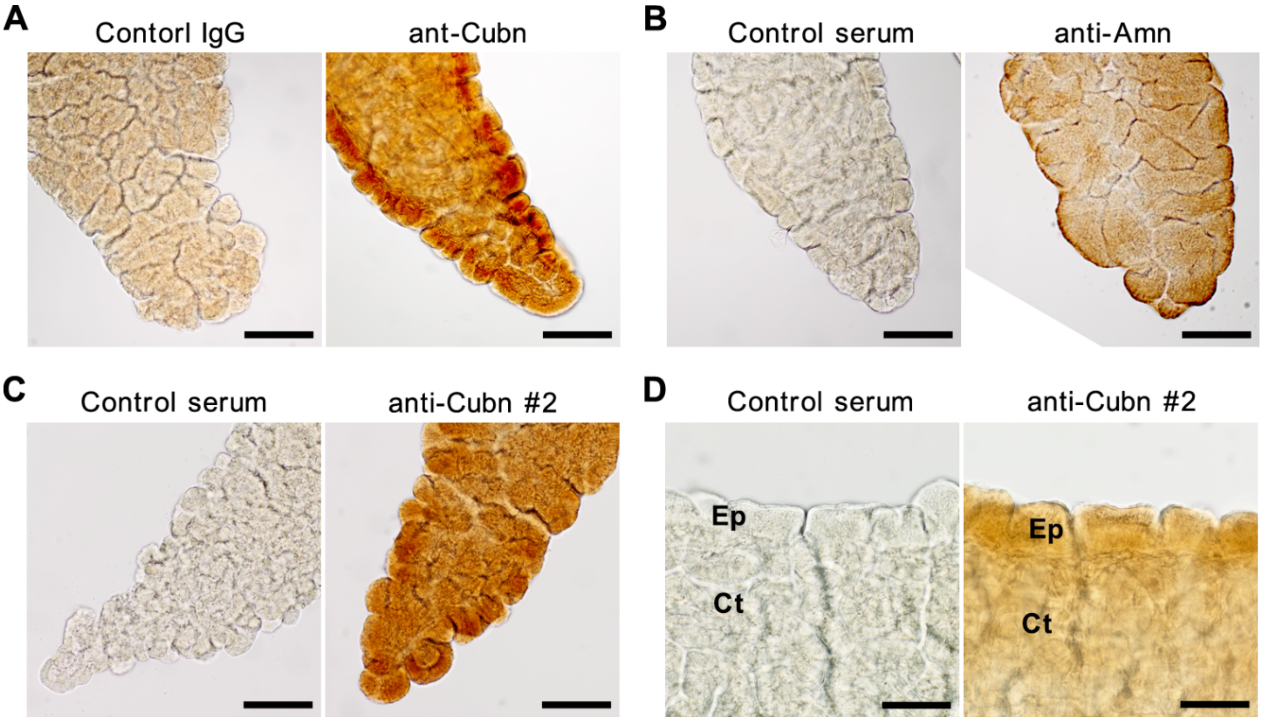


**Figure S2. Immunohistochemistry of Cubn or Amn in the trophotaenia.**

**A-B**. Immunohistochemistry using the Cubn antibody (A) or Amn antiserum (B) in the apical terminal of the trophotaenia. Scale bar: 100 μm. **C-D**. Immunohistochemistry using the Cubn antiserum in the apical terminal (C) or the epithelial layer (enlarged image) (D) of the trophotaenia. Ep, epithelium layer; Ct, connective tissue. Scale bar: 100 μm (C) or 50 μm (D).

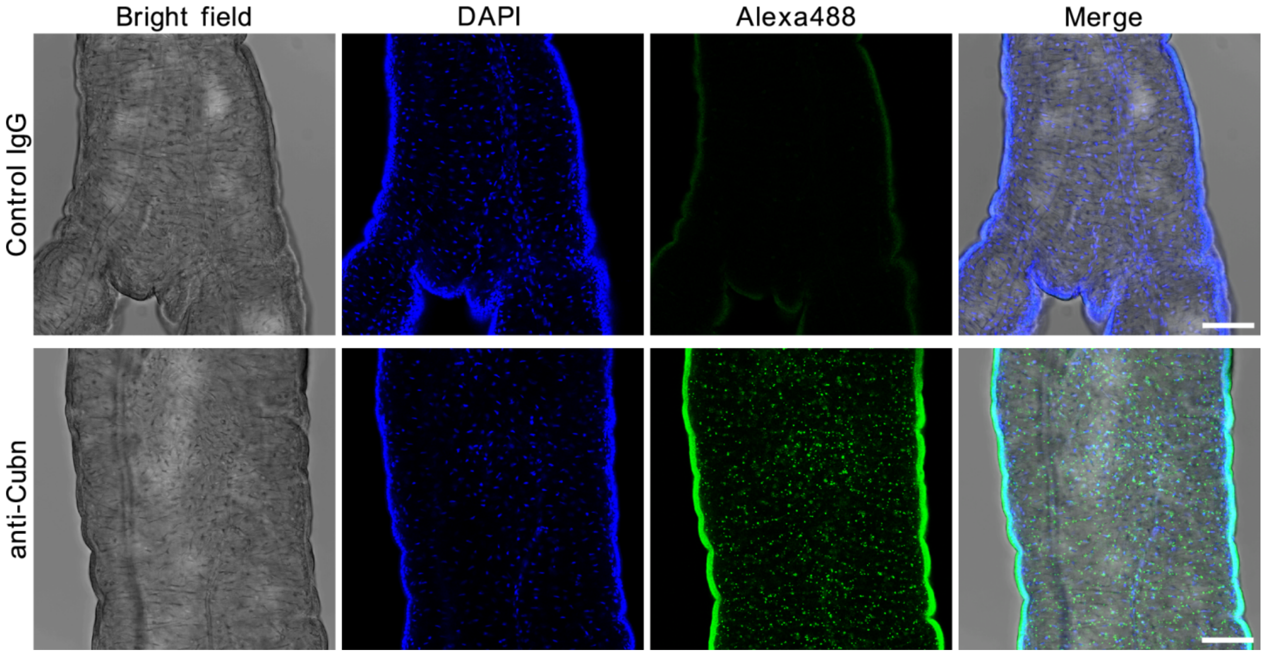


**Figure S3. Fluorescent immunohistochemistry using Cubn antibody in the trophotaenia.**

Confocal microscopy analysis of the trophotaenia of *Xenotoca eiseni* embryos at the fourth week of gestation. Anti-Cubn signals are observed in the epithelial layer cells of the trophotaenia. DAPI, 4′,6-diamidino-2-phenylindole. Scale bar: 20 μm.

```
H.sapiens DAB2 (NP_001334.2) MSNEVETSATN-GQPDQ---------------QAAPKAPSKKEKKKGPEK 34
X.eiseni  dab2               MSTEVENSVPATADPSSPSTASTTISPTTSPVTTTPKAQLKKEKKKVPEK 50
                             **.***.*..  .:*..                ::***  ****** ***

H.sapiens DAB2 (NP_001334.2) TDEYLLARFKGDGVKYKAKLIGIDDVPDARGDKMSQDSMMKLKGMAAAGR 84
X.eiseni  dab2               TDEYLLSRFQGDGVRYKAKLIGIDDVSEARGDKMCQDSMMKLKGMAVAAR 100
                             ******:**:****:***********.:******.***********.*.*

H.sapiens DAB2 (NP_001334.2) SQGQHKQRIWVNISLSGIKIIDEKTGVIEHEHPVNKISFIARDVTDNRAF 134
X.eiseni  dab2               SQGKHKQRIWVNISMSGLKIIDEKSGVIEHEHVVNKISFIARDVTDNRAF 150
                             ***:**********:**:******:******* *****************

H.sapiens DAB2 (NP_001334.2) GYVCGGEGQHQFFAIKTGQQAEPLVVDLKDLFQVIYNVKKKEEEK-KKIE 183
X.eiseni  dab2               GYVCGAEGQHQFFAIKTAQQAEPLVIDLKDLFQVIFNMRKKEAESSQKAE 200
                             *****.***********.*******:*********:*::*** *. :* *

H.sapiens DAB2 (NP_001334.2) EASKAVENGSEALMILDDQTNKLKSGVDQMDLFGDMSTPPDLNSPTESKD 233
X.eiseni  dab2               NGSAVVENGG-ALQSTDGES-KAAQPVEQLDLFGDITTPPDIRAP----- 243
                             :.* .****. **   *.:: *  . *:*:*****::****:.:*

H.sapiens DAB2 (NP_001334.2) ILLVDLNSEIDTNQNSLRENPFLTNGITSCSLPRPTPQASFLPENAFSAN 283
X.eiseni  dab2               --------------------------------------------------

H.sapiens DAB2 (NP_001334.2) LNFFPTPNPDPFRDDPFTQPDQSTPSSFDSLKSPDQKKENSSSSSTPLSN 333
X.eiseni  dab2               ---------------------------------------NSGSS------ 248
                                                                    **.**

H.sapiens DAB2 (NP_001334.2) GPLNGDVDYFGQQFDQISNRTGKQEAQAGPWPFSSSQTQPAVRTQNGVSE 383
X.eiseni  dab2               --------------------------------------------------

H.sapiens DAB2 (NP_001334.2) REQNGFSVKSSPNPFVGSPPKGLSIQNGVKQDLESSVQSSPHDSIAIIPP 433
X.eiseni  dab2               --------------------------------------------------

H.sapiens DAB2 (NP_001334.2) PQSTKPGRGRRTAKSSANDLLASDIFAPPVSEPSGQASPTGQPTALQPNP 483
X.eiseni  dab2               ------------------DLFGTDLFVPPVS------SETSPADLFNNTP 274
                                               **:.:*:*.****      * *. .  :: .*

H.sapiens DAB2 (NP_001334.2) LDLFKTSAPAPVGPLVGLGGVTVTLPQAGPWNTASLVFNQSPSMAPGAMM 533
X.eiseni  dab2               TNTVPSTIPALG--SLQLGPTATSVPAVGMWGTSPAVPAMFP--MPGIVT 320
                              : . :: **     : ** .:.::* .* *.*:. *    *   ** :

H.sapiens DAB2 (NP_001334.2) GGQPSGFSQPVIFGT--SPAVSGWNQPSPFAASTPPPVPVVWGPSASVAP 581
X.eiseni  dab2               PGLRPNFPQPTAFGVPMQPPVWAPQVVPQFSAAPLSPPHLQWGQPATS-- 368
                              *  ..*.**. **.  .*.* . :  . *:*:. .*  : ** .*:

H.sapiens DAB2 (NP_001334.2) NAWSTTSPLGNPFQSNIFPAPAVSTQPPSMHSSLLVTPPQPPPRAGPPKD 631
X.eiseni  dab2               ----------NPFQ-------AMGDHGPSRP------PPRPPVKETPPR- 394
                                       ****       *:. : **        **:** :  **:

H.sapiens DAB2 (NP_001334.2) ISSDAFTALDPLGDKEIKDVKEMFKDFQLRQPPAVPARKGEQTSSGTLS- 680
X.eiseni  dab2               VENSAFTALDPLGDKEKKTGKDMFKNFQIAKLPAIPARKGELMPSSTPPP 444
                             :...************ *  *:***:**: : **:******  .*.* .

H.sapiens DAB2 (NP_001334.2) ------AFASYFNSKVGIPQENADHDDFDANQL-LNKINEPPKPAPRQVS 723
X.eiseni  dab2               ANKESVPFDEYFSNKVGLAQDAADHDDFDINQMSILDGNDAPKQTPVQFT 494
                                   .* .**..***:.*: ******* **: : . *:.** :* *.:

H.sapiens DAB2 (NP_001334.2) LPVTKS------TDNAFENP-----------------FFKDSFGSSQASV 750
X.eiseni  dab2               APAAAPSFTPDLLDAAFSSAPVPNSSAPTLGQDLSHDMFDQAFGAPDPNP 544
                              *.: .       * **...                 :*.::**:.:..

H.sapiens DAB2 (NP_001334.2) ASSQPVSS----------EMYRDPFGNPFA 770
X.eiseni  dab2               FGAPPVAMNTVAQTSGSTDAFGDAFGNPFA 574
                              .: **:           : : *.******
```

**Figure S4. Amino acid sequence alignment for *Homo sapiens* (*H. sapiens*) DAB2 and *Xenotoca eiseni* (*X. eiseni*) dab2-like proteins.**

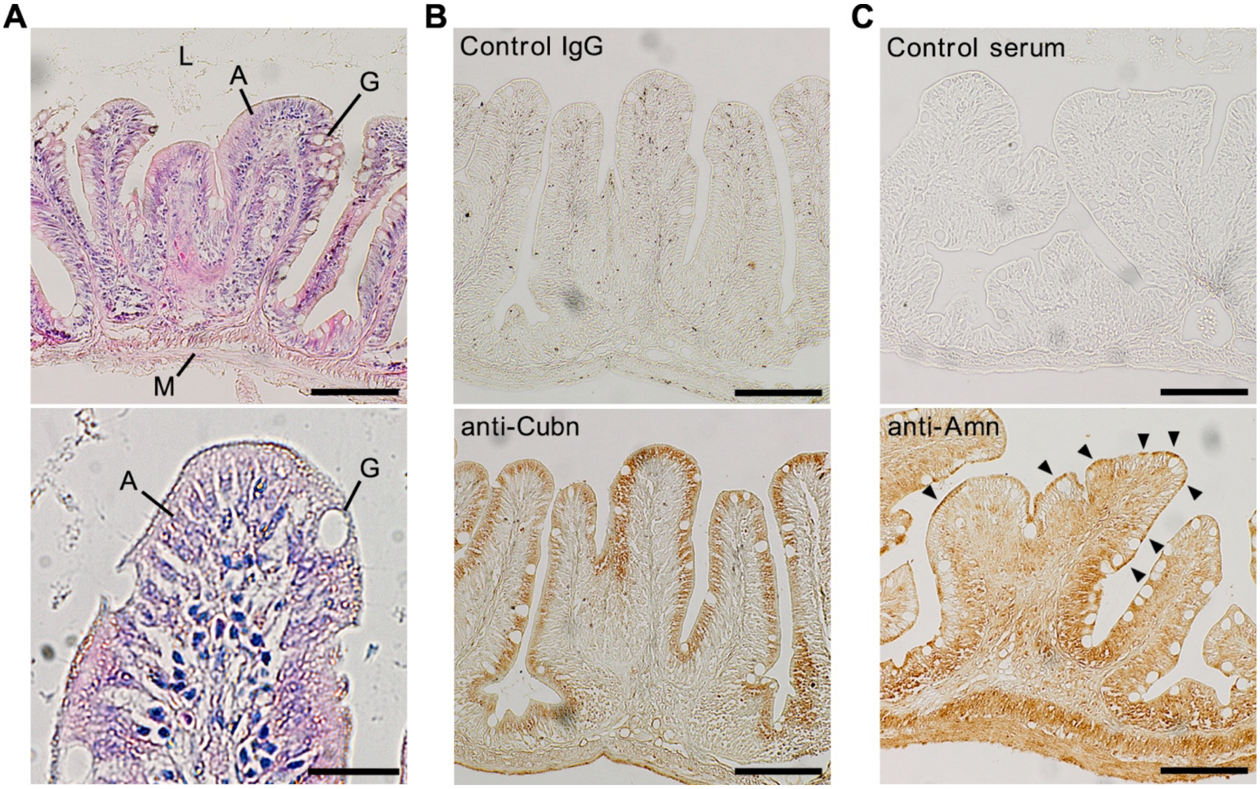


**Figure S5. Immunohistochemistry of Cubn or Amn in the adult intestine of *Xenotoca eiseni*.**

**A**. Hematoxylin and eosin-stained section of the hindgut of an adult female. The lower panel is an enlarged image of an intestinal villus. A, absorbent epithelium; G, goblet cell; L, intestinal lumen; M, muscularis mucosae. Scale bar: 100 μm (upper) and 20 μm (lower). **B**. Immunohistochemistry using the Cubn antibody or control IgG. Scale bar: 100 μm. **C**. Immunohistochemistry using the Amn antiserum or control serum. Arrowheads indicate signal in the apical surface of the absorbent epithelium. Scale bar: 100 μm.

### Additional Tables

| Name | Immunogen | Host | Supplier | ID | Related figures |
|---|---|---|---|---|---|
| CUBN Polyclonal Antibody (anti-Cubn) | Human CUBN | Rabbit | Invitrogen | PA5-83684 | 3C, 3E, 3G, S2, S3 |
| Rabbit IgG Isotype Control (Control IgG) | N/A | Rabbit | Invitrogen | 02-6102 | 3C, 3E, 3G, S2, S3 |
| Cubn antiserum (anti-Cubn #2) | *Xenotoca eiseni* cubn | Mouse | N/A | N/A | S2 |
| Amn antiserum (anti-Amn) | *Xenotoca eiseni* amn | Mouse | N/A | N/A | 3D, S2 |
| Preimmune serum (Control serum) | N/A | Mouse | N/A | N/A | 3D, S2 |
| Goat Anti-Mouse IgG (H+L) Secondary Antibody, Alexa Fluor Plus 488 | Rabbit IgG | Goat | Invitrogen | A32723 | 3E, S3 |
| Horse Anti-Mouse IgG Antibody (H+L), Peroxidase | Rabbit IgG | Horse | VECTOR | PI-2000 | 3D, S2 |
| Goat Anti-Rabbit IgG Antibody (H+L), Peroxidase | Mouse IgG | Goat | VECTOR | PI-1000 | 3C, S2 |
| Goat Anti-Rabbit IgG Antibody (H+L), Biotinylated | Rabbit IgG | Goat | VECTOR | BA-1000 | 3G |

**Table S1. List of antibodies used in the study.**

N/A, not applicable.
